# Phylogenetic diversity efficiently and accurately prioritizes the conservation of aquatic macroinvertebrate communities

**DOI:** 10.1101/709733

**Authors:** Kathryn M. Weglarz, W. Carl Saunders, Andrew Van Wagenen, William D. Pearse

## Abstract

1. Land degradation is a leading cause of biodiversity loss yet its consequences on freshwater ecosystems are poorly understood, exacerbating difficulties with assessing ecosystem quality and the effectiveness of restoration practices.
2. Many monitoring programs rely on macroinvertebrates to assess the biotic effects of degradation and/or restoration and management actions on freshwater ecosystems. The ratio of Observed (O) to Expected (E) macroinvertebrate taxa at a given site—O/E—is often used for this purpose, despite the amount of modeling and data required to generate expectations and difficulties quantitatively assessing the degree of degradation at a site.
3. Despite widespread use in academic biology, phylogenetic diversity is rarely applied in management, regardless of empirical correlations between phylogenetic diversity and management targets such as ecosystem structure and function.
4. We use macroinvertebrate data from 1,400 watersheds to evaluate the potential for phylogenetic metrics to inform evaluations of management practices. These data have been collected since 1998, and have been used to determine the effectiveness of conservation management for the maintenance and restoration of riparian and aquatic systems.
5. Phylogenetic diversity detected degradation as effectively as O/E, despite not having baseline ‘expectation’ data. Site disturbance, road density, and broader environmental drivers such as mean annual temperature strongly predicted site phylogenetic diversity, providing concrete management objectives to increase site health.
6. *Synthesis and applications.* Management efforts targeted solely at taxonomic metrics, such as O/E, have been successfully used to manage sites. We show here that phylogenetic diversity metrics can support such efforts by providing additional information about the kind of species at sites. Given the ease with which such approaches can be applied, we call on others to use them to supplement existing prioritization schemes.

## 1 Introduction

Declines in biodiversity are associated with subsequent loss of ecosystem services (Butchart et al., 2010; Cardinale et al., 2012). This is a particular concern in freshwater ecosystems which are highly vulnerable (Millennium Ecosystem Assessment, 2005). These ecosystems support food web processes and nutrient cycling, purify and supply high quality freshwater resources, regulate sediment and nutrient transport across the landscape, and provide an array of cultural services (Macadam & Stockan, 2015; Vörösmarty et al., 2010). Since the late 1800s, when the link between human health and water quality was formally recognized by the scientific community, freshwater systems have been monitored for conservation and restoration purposes (Bonada, Prat, Resh, & Statzner, 2006). While land degradation is recognized as a leading cause of biodiversity loss (Newbold et al., 2015), its consequences on freshwater ecosystems remain poorly understood (IPBES, 2018). This gap persists in spite of the well-recognized link between land and water management (Bossio, Geheb, & Critchley, 2010), making it difficult to determine the effectiveness of restoration practices (Feld et al., 2011; Melland, Fenton, & Jordan, 2018). To assess the impact of these practices on the biotic community, many monitoring programs survey aquatic macroinvertebrates because these organisms are abundant, provide anywhere from 20-100% of the energy budget to consumers, and are sensitive to changes in water chemistry in predictable ways (Bonada et al., 2006; Macadam & Stockan, 2015). Beyond serving as a biomonitoring tool, macroinvertebrates are integral to decomposition and nutrient cycling and support $31.4 billion in recreational fishing in the US alone (Prather et al., 2013). Unfortunately, the impact of management efforts on freshwater biodiversity, particularly macroinvertebrates and fish, can be difficult to detect owing to the time required for these organisms to return to improved habitats (Meals, Dressing, & Davenport, 2010).

Many monitoring programs rely on macroinvertebrates to assess the biotic effects of degradation because they are relatively easy to collect and identify, and track disruptions more closely than other taxonomic groups (Bonada et al., 2006; Cairns & Pratt, 1993; Friberg et al., 2011). Due to their ubiquity in monitoring, a number metrics have been developed to evaluate the status of macroinvertebrate assemblages (Balloch, Davies, & Jones, 1976; Bonada et al., 2006; Cairns & Pratt, 1993; Hellawell, 2012; Resh & Rosenberg, 1993). Often, these metrics measure ecological structure and are similar to, or based upon, measures of species richness (Yates, Brua, Culp, Chambers, & Wassenaar, 2014). One of the more commonly adopted families of these metrics compares the number of Observed taxa (O) to a statistical Expectation (E) of taxa at a given site (O/E; Hawkins, 2006; Hawkins, Norris, Hogue, & Feminella, 2000; Wright, 1995). This metric became widely applied in management with the development and adoption of the River Invertebrate Prediction and Classification System (RIVPACS) (Wright, Sutcliffe, & Furse, 2000; Wright, 1995), which has since been modified for regions outside of the UK (Hawkins et al., 2000; Moya et al., 2011; Smith, Kay, Edward, Papas, et al., 1999). Such O/E approaches are comparable across systems and relatively easy for managers to interpret (Hawkins, 2006; Wright et al., 2000; Wright, 1995), but, like all approaches, O/E metrics have potential caveats. They are less sensitive to degradation in heterogeneous environments (Hargett, ZumBerge, Hawkins, & Olson, 2007), and uncertainty associated with expectations is often not reflected in the final O/E metric (but see de Zwart, Dyer, Posthuma, & Hawkins, 2006). Further, standards for what constitutes a degraded O/E are often subjective, and statistical deviation from this baseline can be difficult to determine (Hamäläinen, Aroviita, Jyväsjärvi, & Kärkkäinen, 2018). The problem is even more acute given variation (and uncertainty; see above) in expectations of site richness: sites may have the same O/E value even when they have drastically different values of E. For instance, a site with an 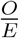 of 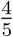 and another with an 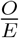 of 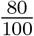 represent the same O/E value of 0.8, but it is unclear whether this value conveys the same information for both sites. Perhaps most critically, however, O/E does not consider the *kinds* of species that are present or absent. The functional diversity of macroinvertebrates reflects an ecosystem’s health and productivity (Schmera, Heino, Podani, Erős, & Dolédec, 2017), and it would be useful, as others have suggested (*e.g.*, Friberg et al., 2011; Yates et al., 2014), for metrics to reflect this.

Even though integrating functional and structural metrics should improve detection of environmental degradation (Yates et al., 2014), macroinvertebrate modeling has been slow to incorporate phylogenetic diversity, which can serve as a proxy for functional diversity (Cavender-Bares, Kozak, Fine, & Kembel, 2009; Faith, 1992; Tucker et al., 2017; Webb, Ackerly, McPeek, & Donoghue, 2002). This is surprising, since much of modern eco-phylogenetic methods were developed from taxonomic metrics developed for macroinvertebrates (reviewed in Pearse, Purvis, Cavender-Bares, and Helmus, 2014; see Izsáki and Papp, 1995; Wright, 1995). Incorporating the evolutionary history of species (phylogeny) into macroinvertebrate monitoring provides the opportunity to incorporate information about the kinds of species within an ecosystem. Under the assumption that more distantly-related species have fewer functional traits in common (but see Mayfield & Levine, 2010), phylogenetic diversity metrics can reveal the structure (Cadotte, Hamilton, & Murray, 2009), stability (Cadotte, Dinnage, & Tilman, 2012; Craven et al., 2018), and primary productivity (Cadotte, Cardinale, & Oakley, 2008; Flynn, Mirotchnick, Jain, Palmer, & Naeem, 2011), of ecosystems while highlighting when certain species loss or gain will have an oversized impact on ecosystem function (Lessard, Fordyce, Gotelli, & Sanders, 2009; Pearse, Chase, et al., 2015; Strauss, Webb, & Salamin, 2006). Yet phylogenetic diversity or evolutionary distinctiveness are rarely incorporated into applied conservation, restoration, and management programs (Cadotte & Tucker, 2018; Díaz et al., 2013; Pearse, Chase, et al., 2015; Tucker et al., 2017), with the notable exception of the EDGE of Existence program (Isaac & Pearse, 2018; Isaac, Turvey, Collen, Waterman, & Baillie, 2007).

The goal of this study is to outline the potential of phylogenetic metrics for macroinvertebrate conservation and monitoring programs. To accomplish this we use a well-established, successful monitoring program as a case study: the PACFISH/INFISH Biological Opinion Effectiveness Monitoring Program (PIBO). PIBO’s goal is to determine the effectiveness of aquatic conservation strategies in riparian and aquatic systems in the Northwestern US, and has sampled stream reaches in over 1,400 watersheds every five years since the late 1990s (see Fig. 1; Henderson, Archer, Bouwes, Coles-Ritchie, & Kershner, 2005). The program’s macroinvertebrate assemblage data provides a direct link between sites’ physical characteristics and the broader biotic community, as macroinvertebrates are prey for the imperiled fishes PIBO was designed to assess. Yet most analyses of the PIBO dataset have focused on physical habitat measures (*e.g.*, Al-Chokhachy et al., 2016; Al-Chokhachy, Roper, Archer, & Miller, 2011; Kershner et al., 2004), and initial attempts to use traditional metrics such as O/E provided insufficient information to evaluate management actions (Irvine et al., 2015). Here, we apply phylogenetic diversity metrics to the macroinvertebrate data within the PIBO dataset to demonstrate their usefulness to biomonitoring programs. We focus on the analysis of three phylogenetic diversity metrics (Faith’s PD, *SES*_*MPD*_, and *SES*_*MNTD*_) as a case study in the potential of phylogenetic diversity metrics to generate actionable management insights.

**Figure 1:**
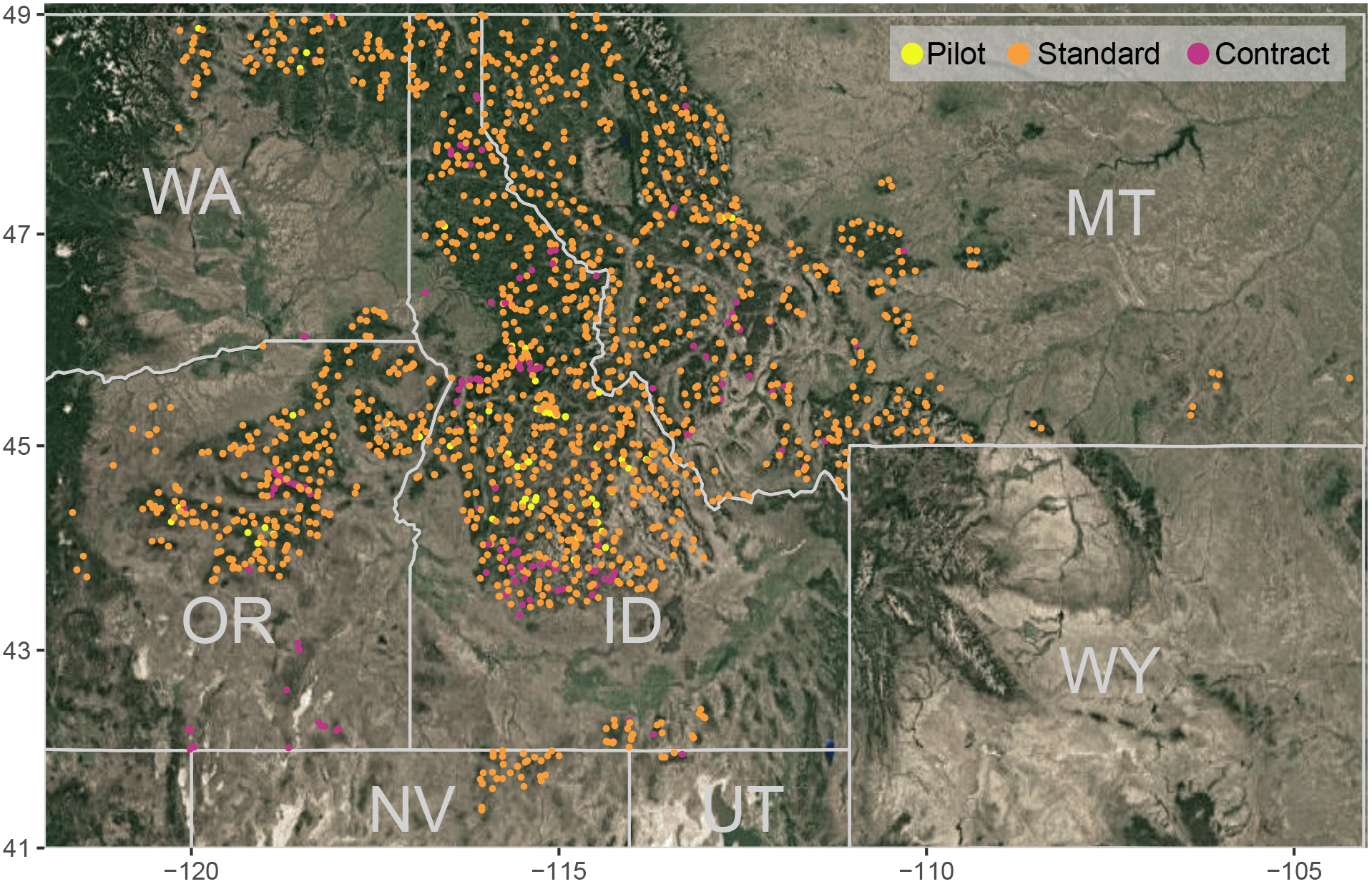
The distribution of the 1667 PIBO sampling sites across the Interior Columbia and Upper Missouri river basins. Sites include 1409 standard sites that are sampled every five years, 100 pilot sites, and 158 contracted or special program sites.

## 2 Methods

### 2.1 Data Collection

The PIBO program collects data from within the Interior Columbia River and Upper Missouri River basins at reach-scale sites (160-400m stream lengths). Sites designated as ‘reference’ (minimally managed) are primarily located in watersheds that have little history of, or no obvious, mining, recent grazing (within 30 years), timber harvest (< 5%), and have minimal road density within the watershed (< 0.5 km/km^2^). Macroinvertebrates are sampled from eight fast-water habitats per site and combined into a composite sample using the protocol recommended by the Center for Monitoring and Assessment of Freshwater Ecosystems at Utah State University (Hawkins et al., 2000). Taxa are identified by the BLM/USU National Aquatic Monitoring Center in Logan (Utah). Physical measures of the environment, such as the frequency of large wood, total dissolved solids, bankfull width, and pool frequency, are sampled concurrently across the PIBO sites. For a complete description of these variables and field methods used, see Henderson et al. (2005). Here we analyze PIBO data collected between 2001 and 2017 across 1667 sites, of which 1477 (88%) were sampled more than once.

### 2.2 Measures of Site Disturbance

All following analyses were conducted in R (3.5.1; R Core Team, 2018), data and code for analyses are available in supplementary materials. To build a macroinvertebrate phylogeny, we searched the TimeTree of Life (Hedges, Marin, Suleski, Paymer, & Kumar, 2015) for species and their congeners, and then added missing species into this phylogeny using dating information from congeners using the ‘bind-replace’ algorithm (*congeneric.merge* in *pez*; Pearse, Cadotte, et al., 2015). The resulting phylogeny contained 164 of the 440 insect taxa in the PIBO data; this was the best coverage we could achieve given many species were identified only to genus (260; and so could not be included in a species-level phylogenetic analysis) and the paucity of genetic data available. The strong performance of our phylogenetic metrics, despite this lack of data, makes our demonstration of the power of phylogenetic metrics, if anything, conservative. However, we note that phylogenetic diversity analyses are resilient to randomly missing species (Isaac & Pearse, 2018). Using this phylogeny, we calculated Faith’s PD (Faith, 1992), *SES*_*MPD*_ (Kembel, 2009; Webb et al., 2002), and *SES*_*MNTD*_ using *pez* (Kembel et al., 2010; Pearse, Cadotte, et al., 2015) under a species-labels null randomization. To compare the relative power of O/E and phylogenetic metrics, we also analyzed the O/E metric values that PIBO calculates for management purposes. PIBO calculates O/E by employing a RIVPACS-type predictive model and comparing the number of taxa expected at similar high quality sites to the number observed at a site (for details see Irvine et al., 2015). This resulted in metrics for 5033 site-year combinations (1477 of the 1667 sites were surveyed more than once) within the PIBO dataset.

### 2.3 Drivers of biotic degradation

To explain potential variation in phylogenetic and O/E metrics, we utilized a range of covariates that describe variation in geoclimatic and anthropogenic drivers across the study area. Table S1 outlines the explanatory variables used in our analysis. We divided these variables into ‘actionable’ variables that management practices may affect, and ‘non-actionable’ which management is unlikely to be able to control. We included the PIBO program’s site designation as ‘managed’ or ‘reference’. To earn the ‘reference’ designation, sites must have no grazing in the watershed (in the last 30 years), <5% of the watershed harvested for timber, no mining (in the last 30 years), and a <0.5 km/km^2^ road density in the watershed. We also included the PIBO program’s condition index of habitat integrity. This condition index is a numeric score ranging from 0 (worst) to 100 (best), calculated by summing independent index values of six variables: residual pool depth, percent pools, diameter of the 50^th^ percentile streambed particle, percent pool tail fines <6mm, large wood frequency, and average bank angle (see Al-Chokhachy, Roper, and Archer (2010) for all details).

With over 40 potential explanatory variables in our dataset, we identified potential drivers of degradation using a two-step approach. First, we used lasso regression (Hastie & Efron, 2013; Tibshirani, 1996) to eliminate explanatory variables with minimal sway on response variables. Lasso regression is a machine learning algorithm designed to reduce a large number of potential explanatory variables to a suite of those that drive a response variable (Hastie & Efron, 2013). With the set of 19 explanatory variables identified by the lasso regression as potentially important for at least one of our four response variables (Faith’s PD, *SES*_*MPD*_, *SES*_*MNTD*_, and O/E), we then used information theoretic criteria to identify the most important predictors of our response variables (Bartoń, 2018). Information theoretic criteria (Burnham & Anderson, 2004) allow for uncertainty in both model specification (which variables are important) and parameters (coefficient estimates) to filter through into model predictions. This allows us to circumvent traditional problems with significance thresholds and their arbitrary decision criteria when dealing with datasets of this size and complexity. We fit mixed effects models using site identity as a random covariate to control for repeated measures from sites through time (Bates, Mächler, Bolker, & Walker, 2015). This ensures that our estimates are not biased by changes in sampling through time and space in the data. Finally, we ensured all explanatory variables were Z-transformed prior to analysis, making each variable’s coefficient a measure of relative importance (Gelman, Carlin, Stern, & Rubin, 2014; Grueber, Nakagawa, Laws, & Jamieson, 2011). This makes variable coefficients directly comparable, such that a variable with a coefficient twice that of another variable is twice as important in driving a response variable, and so allows us to generate more actionable insights.

### 2.4 Identifying sites of concern

To generate site-specific recommendations, we used the random effects of our mixed effect models (Bates et al., 2015). In a mixed effects framework, treating each site identity as a ‘random’ variable controls for uneven sampling of sites while quantifying the impacts of the other explanatory variables. It also generates site-specific estimates for identifying sites with unusually high or low values of our diversity indices (PD, *SES*_*MPD*_, *SES*_*MNTD*_, and O/E) within the context of our model. This method is appropriate for making recommendations within a dataset, but these site-level predictions should not be used to generalize to other nearby sites or to different datasets (as is the case for related random-effects approaches; Hadfield, Wilson, Garant, Sheldon, & Kruuk, 2010; Robinson, 1991). Spatial plotting was conducted using the R packages ‘*rgdal*’, ‘*rgeos*’, ‘*raster*’, and ‘*ggmap*’ (Bivand, Keitt, & Rowlingson, 2018; Bivand & Rundel, 2018; Hijmans, 2018; Kahle & Wickham, 2013).

## 3 Results

The stream-reach condition index that PIBO has been using to quantify management impacts at sampled stream reaches is significantly negatively correlated with *SES*_*MNTD*_ and *SES*_*MPD*_ (the *SES*_*MNTD*_ correlation is strongest; Fig. 2). Thus, sites with more intact aquatic habitat tend to contain macroinvertebrate communities with more close relatives (phylogenetic clustering). Disturbed sites (*i.e*, low index scores), by contrast, are more phylogenetically overdispersed, which is consistent with the pattern that may result from invasion by distantly-related (novel) species at these sites (not explored here, but see Procheş, Wilson, Richardson, & Rejmánek, 2008). Faith’s PD is the only metric we examined that does not incorporate a statistical expectation of diversity, and it is the only metric not to correlate with the condition index (Fig. 2) and for which ‘reference’ sites differed from ‘managed’ sites (Fig. 3).

**Figure 2:**
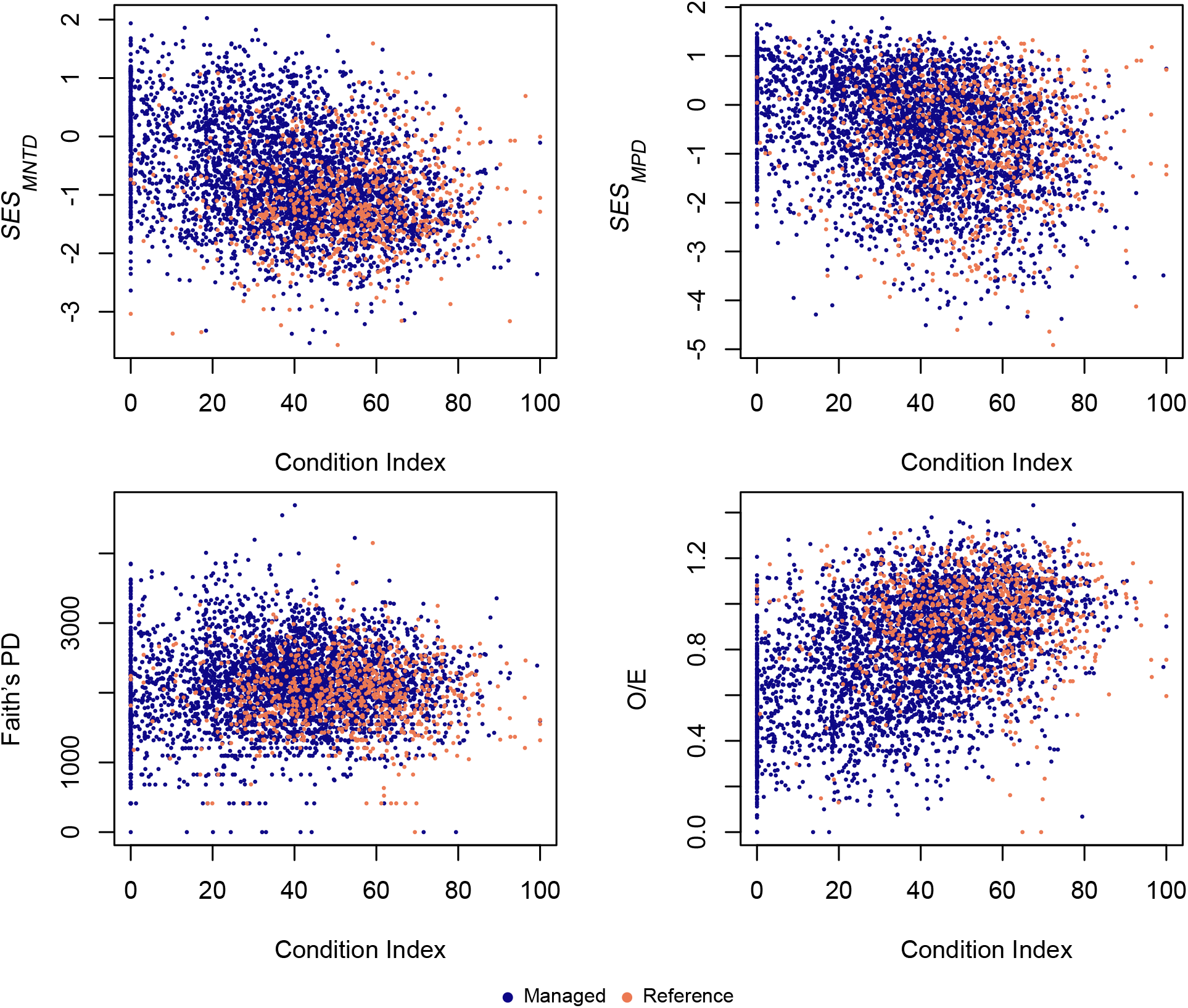
The relationships between condition index and *SES*_*MNTD*_, *SES*_*MPD*_, Faith’s PD, and O/E, showing both managed and reference sites. The results of a linear regression for each of these metrics indicated that site condition index score explained 12.7%, 9.8%, 0.03%, and 18.6% of the variance respectively. Site condition index score was significantly correlated with O/E (*F*_1,4580_ = 1049, *p* < 0.0001, *r*^2^ = 0.186), *SES*_*MPD*_ (*F*_1,4588_ = 504.5, *p* < 0.0001, *r*^2^ = 0.099), and *SES*_*MNTD*_ (*F*_1,4588_ = 641.2, *p* < 0.0001, *r*^2^ = 0.123), but not Faith’s PD (*F*_1,4631_ = 1.51, *p* = 0.220, *r*^2^ = 0.0003). Thus O/E is a better metric of the condition index than our phylogenetic diversity indices, which is to be expected as O/E uses additional data to calculated its expectation, but that its performance is comparable to that of *SES*_*MNTD*_ and *SES*_*MPD*_. However, designation of a site as reference significantly explained only 2.9%, 1.4%, 0.6%, and 5.2% of variance for all metrics respectively (*SES*_*MNTD*_ (*F*_1,4978_ = 141.3, *p* < 0.0001, *r*^2^ = 0.028), *SES*_*MPD*_ (*F*_1,4978_ = 67.44, *p* < 0.0001, *r*^2^ = 0.013), Faith’s PD (*F*_1,5031_ = 33.91, *p* < 0.0001, *r*^2^ = 0.007), and O/E (*F*_1,4975_ = 272.2, *p* < 0.0001, *r*^2^ = 0.052)). This indicates that reference site conditions are not necessarily good benchmarks for macroinvertebrate communities.

**Figure 3:**
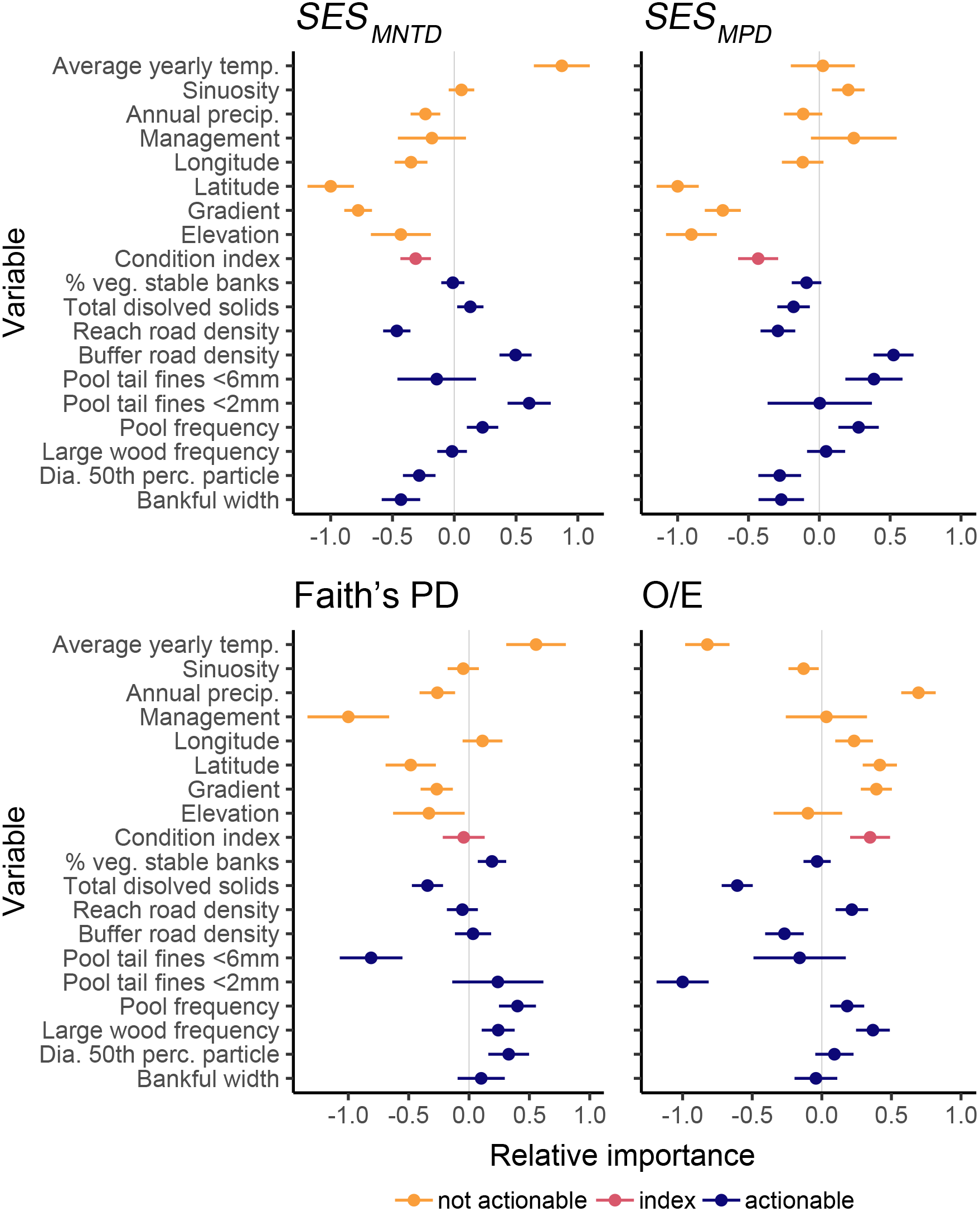
Plots showing the relative importance of each explanatory variable considered in the model for each metric considered. Coefficient values further from zero (either in the positive or negative direction) indicate that the variable has a greater impact on the overall model. For example, the percent of pool tail fines < 2mm present has nearly twice the influence on the *SES*_*MNTD*_ model as pool frequency. Explanatory variables are indicated on the left and are colored by whether or not they are considered potentially actionable by a manager attempting to address habitat degradation. This indicates there are consistently influential actionable explanatory variables.

Many of the important predictors of O/E, *SES*_*MPD*_, and *SES*_*MNTD*_ were the same, while the drivers of PD differed (Fig. 3). Some of the important drivers of *SES*_*MPD*_ and *SES*_*MNTD*_ are not practical for management interventions, such as stream gradient and latitude. There were, however, several potentially actionable factors related to stream geomorphology (average bankfull width, pool tail fines, and number of pools) and road density (in the reach itself and its surroundings). *SES*_*MPD*_ and *SES*_*MNTD*_ differed in the size of pool tail fine sediment that affected them, with *SES*_*MPD*_ more affected by larger particles (< 6mm) than *SES*_*MNTD*_ (< 2mm).

Using our mixed effects framework, we were able to identify sites with unusually high or low (phylogenetic) metric values (Fig. 4). Notably, geographic hot-spots and cold-spots are visible for all variables, which is likely related to the non-actionable diversity drivers we identified, such as latitude.

**Figure 4:**
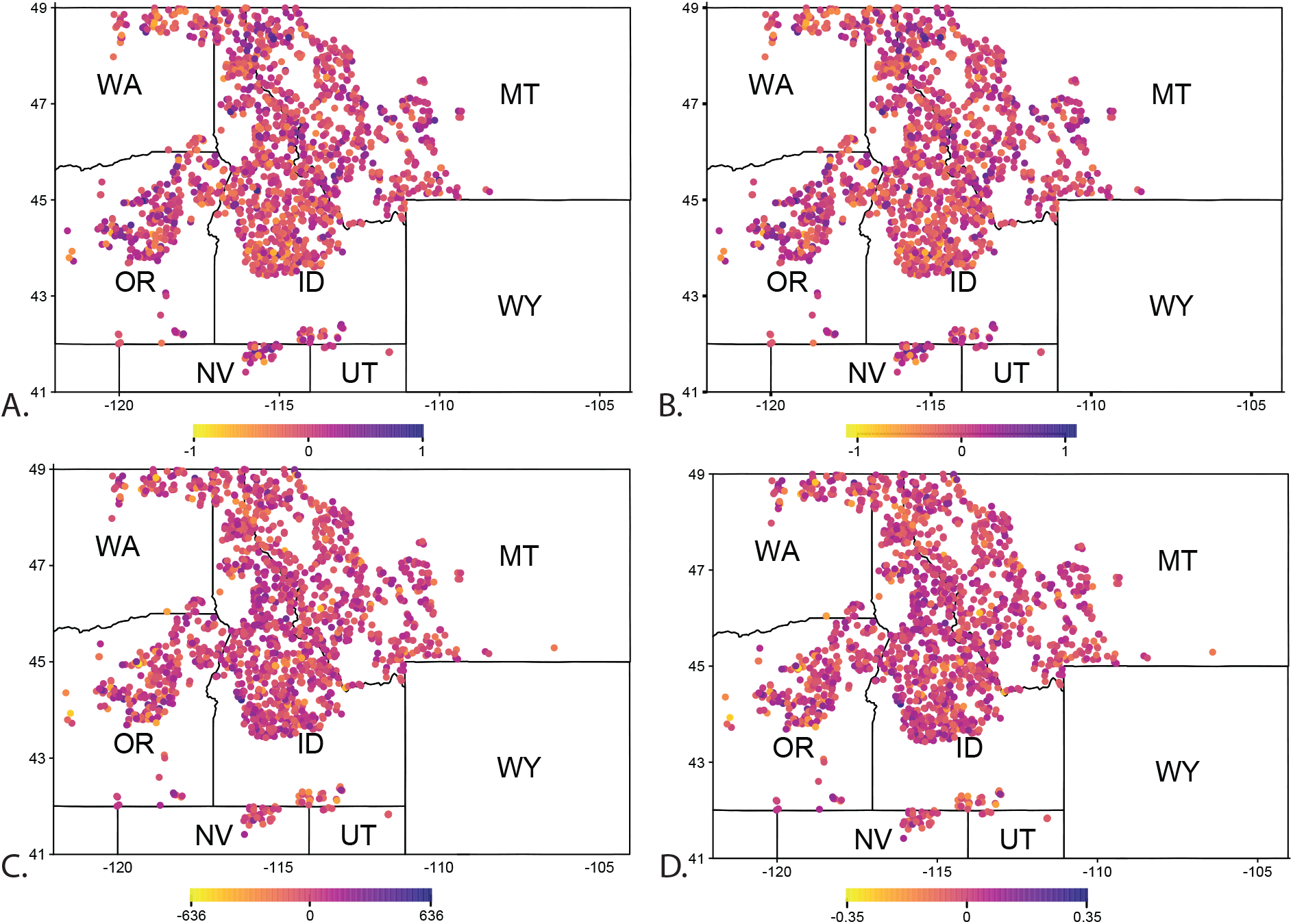
Plots showing site-level variation across each metric, highlighting sites with unusually high or low condition index values given the variation within the dataset for each explanatory variable. In all cases, sites in red should be management priorities. In A (*SES*_*MNTD*_) and B (*SES*_*MPD*_), sites in blue have more positive values, indicating more distantly-related species are found at a site, and values in red have more closely-related species at a site (more negative values), than would be expected given site conditions (management, disturbance, latitude, and all other variables in our models). Again, given clustering (more negative) is found at more disturbed sites, sites in red should be management priorities. In C (Faith’s PD), sites in red have less phylogenetic diversity, and sites in blue more diversity, than would be expected by chance, and thus sites in red are management priorities. Finally, in D (O/E) sites in red have fewer species observed than expected, while sites in blue have more species than the expectation. We emphasize that, as discussed in the text, these estimates should not be used to generalize management decisions to other sites or data, and should only be considered within the context of this modeling exercise.

## 4 Discussion

Assessing and tracking the condition of aquatic and riparian resources on federal lands necessitates tracking changes in sites’ abiotic and biotic conditions to ensure management is effective at achieving its objectives. Many previous analyses of the PIBO dataset use physical habitat measures, compared between reference and managed sites, to assess stream health (e.g. Al-Chokhachy et al., 2016; Al-Chokhachy et al., 2011; Kershner et al., 2004). This work identified important drivers of habitat degradation, but did not quantify the biotic response to such disturbances. Efforts to assess biotic response have primarily focused on the vegetation component of the PIBO dataset, examining the impact of grazing on plants (Coles-Ritchie, Roberts, Kershner, & Henderson, 2007), of plants on stream structure (Hough-Snee, Roper, Wheaton, & Lokteff, 2015; Roper, Jarvis, & Kershner, 2007), and the physical site factors that predict plant invasion (Al-Chokhachy, Ray, Roper, & Archer, 2013; Menuz & Kettenring, 2013). These studies are a solid foundation for understanding the health of these sites, but as a primary prey source for the imperiled fishes PIBO was designed to monitor, an understanding of the macroinvertebrates of the sites is key to achieving management goals. Initial efforts to evaluate the usefulness of macroinvertebrate O/E ratios as the biotic component of conceptual models of degradation failed to find biologically significant results (Irvine et al., 2015), highlighting the need for a more effective metric for measuring the health of macroinvertebrate communities at PIBO sites.

Here we use phylogenetic diversity metrics to quantify the health of macroinvertebrate at sites within the PIBO dataset, demonstrating the continued use of PIBO data to generate actionable insights for land managers. Phylogenetic diversity metrics that incorporate a statistical expectation of diversity (*SES*_*MPD*_ and *SES*_*MNTD*_) are more sensitive than those that do not (Faith’s PD), complementing the current PIBO focus on taxonomic metrics incorporating expectations (O/E; an expectation of taxa that may be present at the site derived through modeling). Below, we discuss the management implications of the drivers we have identified in the data.

### 4.1 Contrasting patterns of phylogenetic and taxonomic diversity

Ratios of Observed to Expected taxa at a site (O/E ratios) are modified measures of species richness that can be compared across sites and studies (Hawkins, 2006). These ratios depend on models of the Probability of taxon Capture (PC) to predict the Expected pool (E), which impacts their performance (Clarke, Wright, & Furse, 2003). O/E ratio performance varies depending on the predictor variables used to generate the expectation and the PC thresholds used to calculate E (Hawkins, 2006; Hawkins et al., 2000). The failure of previous studies to find associations between anthropogenic drivers and the structure of macroinvertebrate assemblages (at PIBO; Irvine et al., 2015), demonstrates how these dependencies limit the application of O/E ratios to locations where PC models have been developed and adequate predictive data can be collected. There are, however, strong conceptual links between the phylogenetic diversity metrics we use here and previous O/E approaches. Faith’s PD augments observed species richness with information about the phylogenetic diversity of those species, and so information about their traits and evolutionary history (Faith, 1992). This added information about the kinds of species within a site is useful, and *SES*_*MPD*_ and *SES*_*MNTD*_ build upon this by incorporating an evolutionary expectation of what a randomly assembled ecological community might resemble (Kembel, 2009; Webb et al., 2002). By combining expectations of community structure with information about the kinds of species in an assemblage, we can pick apart the drivers of community assembly and change. As sites are degraded or restored, it is possible for the number of species within a site to remain constant while the species themselves change; phylogenetic diversity metrics are sensitive to such changes. Critically, in our phylogenetic approach the expected pool comes from the observed data alone: there is no need for comparison sites in order to generate a statistical expectation allowing application of this method in locations that lack PC models. This is a clear practical advantage in favor of the use of phylogenetic diversity metrics.

Phylogenetic diversity is often viewed as a proxy for functional diversity, under the assumption that species that are more distantly related to one another have fewer functional traits in common (Devictor et al., 2010; Mazel et al., 2014; Tucker, Davies, Cadotte, & Pearse, 2018). This is certainly true, at a coarse scale, for macroinvertebrates: species within the same genus often resemble one-another, but many different families and orders are instantly recognizable. Because environmental stressors alter the structure of a community they also predictably alter its phylogenetic diversity (Burns & Strauss, 2011; Cadotte et al., 2008; Cadotte et al., 2012; Flynn et al., 2011; Letcher, 2009). By understanding the phylogenetic structure of healthy communities we can identify damaged communities. Previous work on *Daphnia* indicated that disturbed communities tend to be phylogenetically clustered (Helmus et al., 2010), which is thought to reflect the new environmental filtering pressure of the disturbance. However, disturbed PIBO sites (*i.e.*, those with low condition indices) are instead overdispersed, with high *SES*_*MNTD*_ scores. Undisturbed communities with clustered structures, like PIBO sites, occur in other invertebrate systems such as ants, where invasion results in a reduction in clustering as groups of specialists are lost (Lessard et al., 2009). Here, phylogenetic diversity of macroinvertebrate assemblages is measured within each site (locally) and tends to be clustered in less disturbed sites. Since healthy macroinvertebrate communities are clustered at PIBO sites, we suggest that *SES*_*MPD*_ and *SES*_*MNTD*_ are sensitive enough to pick up on the losses of specialists within these communities. This phylogenetic clustering most likely reflects the high level of habitat heterogeneity found at PIBO sites, with highly specific habitats that host specialists. In contrast, communities at disturbed sites tended to be more over-dispersed – they contain more distantly-related species. This pattern would be consistent with a larger number of invasive (novel, distantly related) or the loss of specialist species, although we do not explicitly test those hypotheses here. Landscape scale stressors that reduce site heterogeneity would result in a loss of close relatives, as they are more likely to be functionally similar (Cavender-Bares et al., 2009). The phylogenetic clustering observed likely further reflects the environmental heterogenity of PIBO sites as assemblages with larger source pools are expected to be more clustered (Cavender-Bares, Keen, & Miles, 2006; Pearse, Jones, & Purvis, 2013; Swenson, Enquist, Pither, Thompson, & Zimmerman, 2006).

### 4.2 Management implications of phylogenetic diversity

Improving and maintaining stream health for macroinvertebrates, and thus the imperiled fishes that prey upon them, depends on identifying the actionable factors that most dramatically disrupt these communities. The relationships between the different physical measures of habitat quality that PIBO collects and the metrics we calculated allow us to highlight factors that are most influential on the macroinvertebrate communities, and some that were surprisingly not. PIBO’s site designation as ‘managed’ or ‘reference’ is not a significant driver of disruption for O/E, *SES*_*MPD*_, or *SES*_*MNTD*_ (Fig. 3, coefficient values overlap 0). During the original study design and implementation phase of the PIBO program, reference sites were selected to encompass both the greatest spatial extent possible as well as the range of variation in habitat conditions resulting from natural disturbances (*e.g.*, wildlife, landslides). Furthermore, not all anthropogenic disturbances (*e.g.*, recreation) were considered in the initial selection of reference sites. As a result, the binary classification of managed/reference alone is apparently insufficient to distinguish between intact and impaired macroinvertebrate assemblages. In contrast, quantitative measures of sites’ abiotic condition (in the form of actionable explanatory variables) provide clear evidence that management affects these drivers of macroinvertebrate assemblage. PIBO’s overall abiotic site condition index, significantly correlates with O/E, *SES*_*MPD*_, and *SES*_*MNTD*_, but not Faith’s PD. This metric is the only of the four diversity metrics that does not incorporate some expectation of diversity. This result demonstrates that a statistical expectation of diversity is a useful component for metrics and is consistent with results from previous studies of the PIBO dataset. Other abiotic factors such as roads in a reach and/or its surroundings are some of the strongest factors influencing community structure. Road construction increases fine sediment and adversely impacts macroinvertebrates (Wood & Armitage, 1997). Measures of fine sediment, specifically percent pool tail fines present, also reflect this relationship, driving overdispersion nearly as strongly, or in the case of *SES*_*MNTD*_ more so than, roads. Because *SES*_*MPD*_ reflects broader patterns across the entire phylogeny, this is consistent with habitat filtering of entire lineages on the basis of the percent of pool tail fines <6mm present, and the percent of pool tail fines <2mm present driving competition among close relatives, thus having a greater impact on *SES*_*MNTD*_. The bankfull width of the reach and frequency of pools are about half as influential as the previously mentioned measures, indicating that overall stream structure has some impact on the communities present. The amount of Total Dissolved Solids (TDS) and percent vegetatively stable banks have a comparatively minor influence on community structure, supporting previous findings that these measures may have little impact on macroinvertebrates and should not be the focus of management actions (Mazeika, Sullivan, Watzin, & Hession, 2004; Timpano, Schoenholtz, Zipper, & Soucek, 2010). That TDS were not important in our model does not mean TDS do not influence macroinvertebrates (Clements & Kotalik, 2016), particularly as measurement of this variable may be limited by measuring at base flows (Cey, Rudolph, Parkin, & Aravena, 1998). Rather, measures of pool tail fines and roads appear to be better indicators of macroinvertebrate assemblage response to management activities within the PIBO dataset.

Surprisingly, the frequency of large wood had little influence on macroinvertebrate communities when other explanatory variables such as sedimentation are included. Large wood has been hypothesized to be an important moderator of human stressors, specifically sedimentation (Irvine et al., 2015), but does not appear to strongly impact assemblage structure in these data. This indicates that indirect effects of large wood have mostly been accounted for by other explanatory variables in our model. In general, these results suggest that stream macroinvertebrates are consistently sensitive to road density, overall stream structure, and sedimentation. To remediate degraded macroinvertebrate assemblages, managers should focus efforts on actionable factors, such as roads, fine sediment, and the number of pools per kilometer, that have a large influence on community structure.

Simply identifying overdispersed communities does not adequately prioritize sites for management purposes. Community response to disturbance must be driven by physical measures that are ‘actionable’ and thus may be affected by management practices. Our statistical models identified a number of drivers of diversity in the PIBO data that are not likely to be influenced by management actions but are still important determinants of community structure (see Fig. 3). Of these non-actionable variables, stream gradient, elevation, and latitude are important predictors of *SES*_*MNTD*_ and *SES*_*MPD*_, and geographic ‘hotspots’ and ‘coldspots’ are visible for all variables in Figure 4. These variables have nearly double the influence on our models as both fine sediment and road density, reflecting latitudinal gradient in biodiversity (Hillebrand, 2004), aquatic insect sensitivity to temperature (Johnson & Jones, 2000), and known patterns of decreased diversity in aquatic communities at higher elevations (Altermatt, Seymour, & Martinez, 2013). Even though these variables strongly influence the model, they are a result of site location and thus cannot be addressed by land managers. We suggest that, all other things being equal, those sites that are strongly impacted by these factors (*i.e.*, are at high latitudes and are hot), should be a lower management priority. Some managers may be able to address factors that we designate as non-actionable. For example, grazing is known to impact instream temperatures (Kovach, Muhlfeld, Al-Chokhachy, Ojala, & Archer, 2019), and thus altering grazing intensity could be used to alter temperature and thus help remediate the site. By accounting for what cannot be changed and acting on what can, managers will effectively and efficiently be able to apply these metrics.

## 5 Conclusion

O/E ratios has provided strong management insight for decades, but like all approaches it has strengths and disadvantages. Here we have shown how phylogenetic diversity metrics, which have less ecological data requirements, can provide meaningful insights into the kinds of species present in communities. In this system, we have found phylogenetic diversity, when combined with an analysis targeted at quantifying the potential benefits of management interventions, to be of great insight. We encourage others to experiment with these new metrics and approaches, in the hope that they will be of use in other monitoring programs of macroinvertebrates and other taxa.

## Author’s Contributions

WDP, WCS, and KMW conceived the ideas and designed methodology; WCS and AVW collected, managed, and provided insight about the data; KMW and WDP analysed the data; KMW and WDP led the writing of the manuscript. All authors contributed critically to the drafts and gave final approval for publication.

## Acknowledgments

We are grateful to anonymous reviewers, and the editorial board, for their help improving this manuscript. WDP, KMW, and the Pearse lab are funded by NSF ABI-1759965, NSF EF-1802605, and USDA Forest Service agreement 18-CS-11046000-041.

## Data accessibility

Data are publicly available by request through the USDA Forest Service PACFISH/INFISH Biological Opinion Monitoring Program (https://www.fs.usda.gov/detail/r4/landmanagement/resourcemanagement/ and the code to conduct analyses is available in the supplemental information.

