## Supplemental Table 1 for "Phylogenetic diversity efficiently and accurately prioritizes the conservation of aquatic macroinvertebrate communities"

### Supplemental Tables

Table 1: Definitions of all variables considered for inclusion in complete model with information on which variables made it through the lasso regression and into the AIC analysis. Additionally, if a land manager could potentially take action to improve the measurement of that variable they are designated 'Actionable'; for example, site latitude may correlate with diversity, but cannot be altered by management practices at a site. Some non-actionable variables may be altered by management practices, but only on a site-specific basis. For example, temperature may be altered by a change in grazing regimes, but this is only possible if that site is grazed.

| **Variable** | **Definition** | **Actionable?** | **AIC?** |
|---|---|---|---|
| Year | The year sampling occurred. | No | No |
| Latitude | Latitude coordinates in Albers Equal Area. | No | Yes |
| Longitude | Longitude coordinates in Albers Equal Area. | No | Yes |
| Elevation | Elevation in feet above sea level. | No | Yes |
| Management designation | Land management types derived by PIBO: Reference (aka minimally managed) or Managed. | No | Yes |
| Stream flow | Categorical call of reach stream flow at time of sample. | Yes | Yes |
| Condition index | Numeric score 0 (worst) - 100 (best) ranking habitat integrity. Index score is calculated by summing values of residual pool depth, percent pools, diameter of 50th per. particle, percent pool tail fines <6mm, large wood frequency, and average bank angle, and scaling 0 - 100. | Index | Yes |
| Total dissolved solids | Measure of the concentration of ionized materials in water, or the ability of water to conduct electrical current. | Yes | Yes |
| Average bankfull width from transects | The average of the bankfull widths at the 21-25 transects measured at each reach. | Yes | No |
| Reach length | Length of sampling reach measured along the thalweg. | No | No |
| Gradient | The difference between the elevation of the water surface at the bottom of the reach and the elevation of the water surface at the top of the reach divided by the reach length. | No | Yes |

Table 1: continued from previous page

| **Variable** | **Definition** | **Actionable?** | **AIC?** |
|---|---|---|---|
| Sinuosity | Reach length divided by the straight valley length from the bottom of the reach to the top of the reach. | No | Yes |
| Residual pool depth | Average of the residual pool depth values for all pools in a reach, which are calculated by subtracting pool tail depth from max depth. | Yes | No |
| Pool frequency | Number of pools within the sampled reach standardized to pools per km. | Yes | Yes |
| Pool percentage | Sum of all qualifying pool lengths divided by the reach length, multiplied by 100. | Yes | No |
| Bankfull width-to-depth ratio at transects | Average of the bankfull width-to-depth ratio from 10 cross sections, calculated as bankfull width divided by the bankfull depth. | Yes | Yes |
| Wetted width-to-depth ratio at transects | Average of the wetted width-to-depth ratio from 10 cross sections, calculated as wetted width divided by the wetted depth. | Yes | No |
| Diameter of $50^{th}$ percentile streambed particle | 100 particles are measured per reach, with five particles collected along each transect. | Yes | Yes |
| Pool tail fines <2mm | The percentage of particles <2mm calculated three times using a 0.36m x 0.36m grid with 50 intersections and averaged for each pool, then averaged for all pools within the reach. | Yes | Yes |
| Pool tail fines <6mm | The percentage of particles <6mm calculated three times using a 0.36m x 0.36m grid with 50 intersections and averaged for each pool, then averaged for all pools within the reach. | Yes | Yes |
| Percent stable banks | The number of covered stable, uncovered stable, and false bank measurements divided by the total number of measurements and multiplied by 100. | Yes | No |
| Percent vegitatively stable banks | The number of covered stable and false bank measurements divided by the total number of measurements and multiplied by 100. | Yes | Yes |
| Under cut percentage | Number of transects with bank angles <90 degrees divided by the total number of transect bank measurements and multiplied by 100. | Yes | No |

Table 1: continued from previous page

| **Variable** | **Definition** | **Actionable?** | **AIC?** |
|---|---|---|---|
| Bank angle | Average of all bank angle measurements with bank angles less than 45 degrees summarized as 45 degrees. | Yes | No |
| Large wood frequency | Number of wood pieces with length ≥1m and diameter ≥0.1m within the reach and standardized to per kilometer. | Yes | Yes |
| Large wood volume | Volume of wood pieces with length ≥1m and diameter ≥0.1m measured within the reach and then standardized to per kilometer. | Yes | No |
| Buffer road density | The sum length of all roads in a given buffer divided by the area in square kilometers of the same buffer. | Yes | Yes |
| Catchement road density | The sum length of all roads in a given catchement divided by the area in square kilometers of the same catchement. | Yes | No |
| Reach road density | The sum length of all roads in a given reach divided by the area in square kilometers of the same reach. | Yes | Yes |
| Segment road density | The sum length of all roads in a given segment divided by the area in square kilometers of the same segment. | Yes | No |
| Annual precipitation | Annual total precipitation (rain and melted snow). | No | Yes |
| Average yearly temperature | Average air, not in-stream, temperature in a given catchmentfor an entire given year. | No | Yes |
| Ecoregion III designation | Level III mapping describes small ecological areas nested within level II regions. | No | No |
| Ecoregion IV designation | Level IV mapping describes small ecological areas nested within level III regions, which are nested within the still larger level II. | No | No |
| Percent burned in segment | Percent of segment burned over a five-year period, derived from the geoprocessing of the LandFire, satellite data provided by a USGS, US Forest Service and BLM created, disturbance dataset. | Yes | No |

Table 1: continued from previous page

| **Variable** | **Definition** | **Actionable?** | **AIC?** |
|---|---|---|---|
| Percent burned in catchement | Percent of catchement burned over a five-year period, derived from the geoprocessing of the LandFire, satellite data provided by a USGS, US Forest Service and BLM created, disturbance dataset. | Yes | No |
| Percent burned in reach | Percent of reach burned over a five-year period, derived from the geoprocessing of the LandFire, satellite data provided by a USGS, US Forest Service and BLM created, disturbance dataset. | Yes | No |
| Percent burned in buffer | Percent of buffer burned over a five-year period, derived from the geoprocessing of the LandFire, satellite data provided by a USGS, US Forest Service and BLM created, disturbance dataset. | Yes | No |
| Average yearly max. temp. | Maximum air temperature in a given catchment for an entire given year. | No | No |
| Average yearly min. temp. | Minimum air temperature in a given catchment for an entire given year. | No | No |
